# Regulation of CHD2 expression by the *Chaserr* long noncoding RNA is essential for viability

**DOI:** 10.1101/536771

**Authors:** Aviv Rom, Liliya Melamed, Micah Jonathan Goldrich, Rotem Kadir, Matan Golan, Inbal Biton, Rotem Ben-Tov Perry, Igor Ulitsky

## Abstract

Genomic loci adjacent to genes encoding for transcription factors and chromatin remodelers are enriched for long non-coding RNAs (lncRNAs), but the functional importance of this enrichment is largely unclear. Chromodomain helicase DNA binding protein 2 (*Chd2*) is a chromatin remodeller with various reported functions in cell differentiation and DNA damage response. Heterozygous mutations in human *CHD2* have been implicated in epilepsy, neurodevelopmental delay, and intellectual disability. Here we show that *Chaserr*, a highly conserved lncRNA transcribed from a region near the transcription start site of *Chd2* and on the same strand, acts in concert with the CHD2 protein to maintain proper *Chd2* expression levels. Loss of *Chaserr* in mice leads to early postnatal lethality in homozygous mice, and severe growth retardation in heterozygotes. Mechanistically, loss of *Chaserr* leads to substantially increased *Chd2* mRNA and protein levels, which in turn lead to increased transcriptional interference by inhibiting promoters found downstream of highly expressed genes. We further show that *Chaserr* production represses *Chd2* expression solely in *cis*, and that the phenotypic consequences of *Chaserr* loss are rescued when *Chd2* is perturbed as well. Targeting *Chaserr* is thus a potentially viable strategy for increasing CHD2 levels in haploinsufficient individuals.

## Introduction

The mammalian transcriptome is highly complex, and contains tens of thousands of non-coding RNA genes^1–4^. A significant subset of these, referred to as long noncoding RNAs (lncRNAs), are at least 200 nucleotides (nt) in length, 3’ polyadenylated, and 5’ capped, and are therefore structurally similar to mRNAs, but lack protein coding potential^5^. Only a small portion of lncRNAs have been functionally characterized, and only very few of these have been studied in the context of organismal development^6^. Multiple lines of evidence point to a strong link between lncRNA functions and those of chromatin modifying complexes^7,8^. Numerous chromatin modifiers have been reported to interact with lncRNAs^8^. In addition, lncRNAs in vertebrate genomes are enriched in the vicinity of genes that encode for transcription-related factors^2,9^, including numerous chromatin-associated proteins, but the functions of the vast majority of these lncRNAs remain unknown.

We hypothesized that the proximity of some lncRNA genes to genes involved in chromatin biology may imply a functional connection. To explore the biology of such interactions, we focused on one of the most conserved lncRNAs in vertebrates, found in close proximity to *Chromodomain Helicase DNA Binding Protein 2* (*Chd2*). CHD2 is an ATP-dependent chromatin-remodeling enzyme, which together with CHD1 belongs to subfamily I of the chromodomain helicase DNA-binding (CHD) protein family. Members of this subfamily are characterized by two chromodomains located in the N-terminal region and a centrally located SNF2-like ATPase domain^10^, and facilitate disassembly, eviction, sliding, and spacing of nucleosomes^11^. There are conflicting reports on the genomic occupancy of CHD2. According to one report, based on ChIP-seq data obtained with antibodies against CHD1 and CHD2, both proteins bind predominantly in the proximity of gene promoters and share up to 60% of their DNA binding sites in human cell lines^12^. Another study used MNase-ChIP-seq of endogenously tagged *Chd1* and *Chd2* in mouse embryonic stem cells (mESCs)^13^, and reported that the two proteins have different binding patterns – CHD1 binds predominantly to promoter regions, whereas CHD2 is associated with gene bodies of actively transcribed genes. CHD2 has also been linked to the deposition and incorporation of the H3.3 histone variant at transcriptionally activated genes^12,14,15^ and at DNA damage foci, with the latter activity promoting repair of double strand breaks^16^.

Mice homozygous for a gene-trap stop cassette in intron 27 of *Chd2* survive until E18.5 with a marked growth retardation, and no viable offspring of these mice can be recovered^17^. Heterozygotes with this mutation show increased mortality at postnatal days 1–4, and in the long-term they exhibit growth retardation, shorter life spans, and altered morphology in various organs. However, a dominant negative effect of the truncated protein could not be excluded in this model. A different model for *Chd2* loss-of-function was recently created by the International Mouse Phenotyping Consortium, where exon 3 was replaced by a lacZ cassette and a stop signal^18^. No significant changes in mortality and aging were reported for these mice, but they exhibit slightly decreased body weight and length, skeletal abnormalities, an abnormal bone structure, decreased fat amount and bone mineral density, and abnormalities in blood composition, such as decreased erythrocyte cell number, hemoglobin content, and mean platelet volume (http://www.mousephenotype.org/).

In humans, *CHD2* haploinsufficiency is associated with neurodevelopmental delay, intellectual disability, epilepsy, and behavioral problems (reviewed in^19^). Studies in mouse models and cell lines also implicate *Chd2* in neuronal dysfunction: perturbations of *Chd2* affect neurogenesis in the mouse developing cerebral cortex^20^ and in human stem cells differentiated to neurons^21^, and loss of a single *Chd2* copy leads to deficits in neuron proliferation and a shift in neuronal excitability^22^. Approaches for increasing CHD2 levels may thus have therapeutic relevance.

## Results

LncRNAs are spread throughout vertebrate genomes, but have a notable enrichment in proximity to chromatin- and transcription-related genes^2,9^. One such lncRNA, annotated as *1810026B05Rik* in mouse (which we denote as *Chaserr*, for *CHD2 adjacent, suppressive regulatory RNA*) and *LINC01578*/*LOC100507217* in human (*CHASERR*), is an almost completely uncharacterized lncRNA, found upstream of and transcribed from the same strand as *Chd2* (**Fig. 1a**). *Chaserr* has five exons, is polyadenylated, and is a bona fide lncRNA according to PhyloCSF^23^ (**Extended Data Fig. 1a**), CPAT^24^ (coding probability 0.296), and CPC^25^ (coding potential score −1.23). According to FANTOM5 transcription start site (TSS) annotations, 3P-seq poly(A) site mapping, and RNA-seq data, *Chaserr* transcript is independent of *Chd2* (**Fig. 1a**). The ‘tandem’ organization with *Chd2*, *Chaserr* exon-intron structure, and parts of *Chaserr* sequence are conserved throughout vertebrates (**Fig. 1a**), which makes it one of the most conserved mammalian lncRNAs^9,26^. According to RefSeq annotation, the last exon of *Chaserr* in mouse overlaps *Chd2*; however, according to RNA-seq and 3P-seq data from various tissues, the predominant *Chaserr* isoform ends ~500 bp after its last 3’ splice site, ~2.2 Kb upstream of the *Chd2* TSS (as in GENCODE transcript ENSMUST00000184554), and we therefore considered this isoform in further studies. The RNA product of *Chaserr* is localized in the nucleus, in proximity to the *Chd2* site of transcription (**Fig. 1b**). This nuclear enrichment is due at least in part to nonsense mediated decay (NMD) that acts on *Chaserr*, likely triggered by a 117 codon non-conserved ORF that ends in the 2^nd^ exon (**Extended Data Fig. 1b-c**).

**Figure 1.**
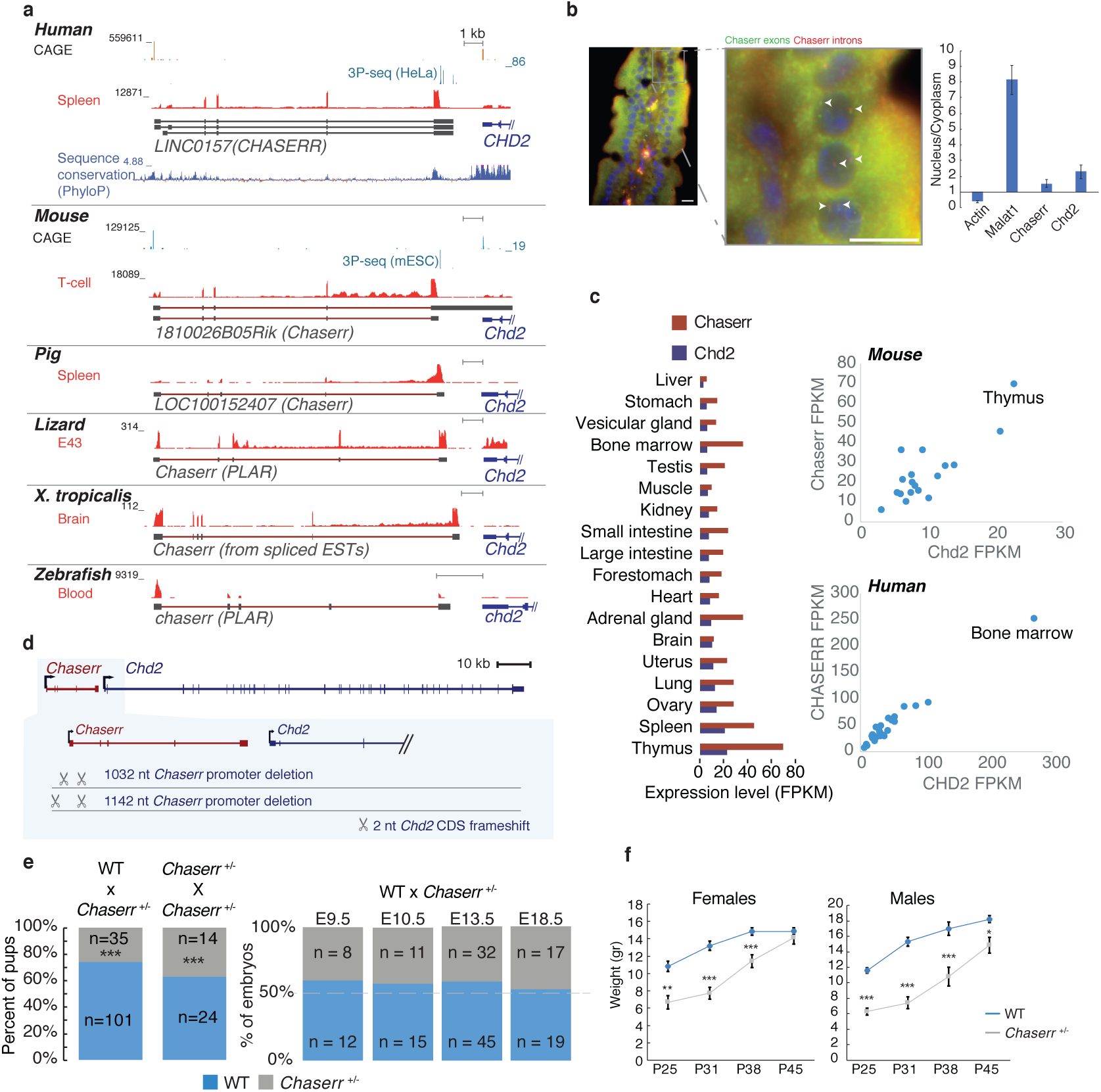
*Chaserr* is essential for postnatal viability. **a,** Genomic locus of *Chaserr* in the indicated vertebrate genomes. CAGE read coverage is from the FANTOM5 project ^64^. 3P-seq data are from ^65^. RNA-seq data are from HPA^66^, ENCODE (mouse), FAANG (pig), SRP009831 (lizard), SRP039546 (xenopus), SRP024369 (zebrafish). **b,** Left: Single molecule FISH with probes targeting *Chaserr* introns (red) and exons (green) in mouse duodenum. Co-localization of introns and exons signal marked by white arrowhead. Scale bar shows 10µm. Right: qRT-PCR comparing cytoplasm and nuclear fractions in mESCs. **c,** Left: *Chaserr* and *Chd2* RNA expression across different tissues profiled in the mouse BodyMap^67^. Right: Scatter plots for *Chaserr* and *Chd2* expression in mouse BodyMap and human HPA^66^ projects. **d,** Schematic of the positions of regions targeted by gRNAs used to generate *Chaserr*^−/−^ and *Chd2*^m/m^ mice. **e,** Observed rates of *Chaserr*^+/–^ and *Chaserr^−/−^* neonates. **f,** Weights of neonates at the indicated age (males n=7–15, females n=6–18 per group).Error bars show S.E.M. *P<0.05 **P<0.01 and ***P<0.001 (two-sided t-test).

*Chaserr* and *Chd2* mRNA are tightly co-expressed across a panel of human and mouse tissues, and during mouse development according to ENCODE and FANTOM5 data **(Fig. 1c** and **Extended Data Fig. 1d-e)**. Interestingly, both genes are expressed at appreciable levels in all studied samples, with particularly high expression in lymphocytes. The ratio between the expression levels of the two genes is also similar across the tissues, with notable exceptions of neuronal cells and fibroblasts, where *Chaserr* levels are relatively high (**Fig. 1c and Extended Data Fig. 1d-e**).

In order to study its function, we generated *Chaserr* null alleles in mice by injection of CRISPR/Cas9 with a pair of gRNAs targeting sequences flanking the promoter and the first exon of *Chaserr* (**Fig. 1d**). This resulted in two different *Chaserr*^+/–^ mouse strains that carry deletions of 1032 and 1142 bps (**Extended Data Fig. 1f**), which were sufficient to abolish expression of mature *Chaserr* (see below), and the two lines were phenotypically indistinguishable from each other, and so we used them interchangeably in subsequent experiments. Strikingly, out of 38 pups born following crosses between *Chaserr*^+/–^ mice, we observed no *Chaserr*^−/−^ pups. The numbers of *Chaserr*^+/–^ pups at weaning also deviated from expected Mendelian ratios (~37%, P < 10^−8^, **Fig. 1e**), and the surviving mice exhibited substantial growth retardation, occasional malocclusion, and neonatal lethality (**Fig. 1f** and **Extended Data Fig. 1g-i**). Pathological analysis of *Chaserr*^+/–^ mice revealed a wide range of abnormalities, including fat depletion, kyphosis, and thymic depletion, neither of which was highly penetrant. *Chaserr*^+/–^ females very rarely became pregnant, therefore we focused on crosses between *Chaserr*^+/+^ females and *Chaserr*^+/–^ males for most of this study. Out of 136 pups born from such crosses, 35 were *Chaserr*^+/–^ (~25%), significantly less than the expected 50% (P < 10^−5^, **Fig. 1e**), suggesting two copies of *Chaserr* are required for proper postnatal survival. In contrast, embryos at four different developmental stages up to E18.5 were recovered with the expected ~50% *Chaserr*^+/–^ vs. *Chaserr*^+/+^ ratios, suggesting that a single copy of *Chaserr* is sufficient for viable embryonic development (**Fig. 1e**).

Due to the close proximity and co-expression of *Chaserr* and *Chd2*, we tested whether *Chd2* expression is affected in *Chaserr*^+/–^ embryos. *Chd2* mRNA was significantly upregulated by ~1.5-fold during embryonic development at the examined time points (E9.5, E10.5, and E13.5), in mouse embryonic fibroblasts (mEFs) derived from E13.5 embryos, and in four adult tissues (**Fig. 2a)**. Western blot (WB) in E13.5 embryos and in the adult tissues also demonstrated strong up-regulation of CHD2 protein (**Fig. 2b-c**).

**Figure 2.**
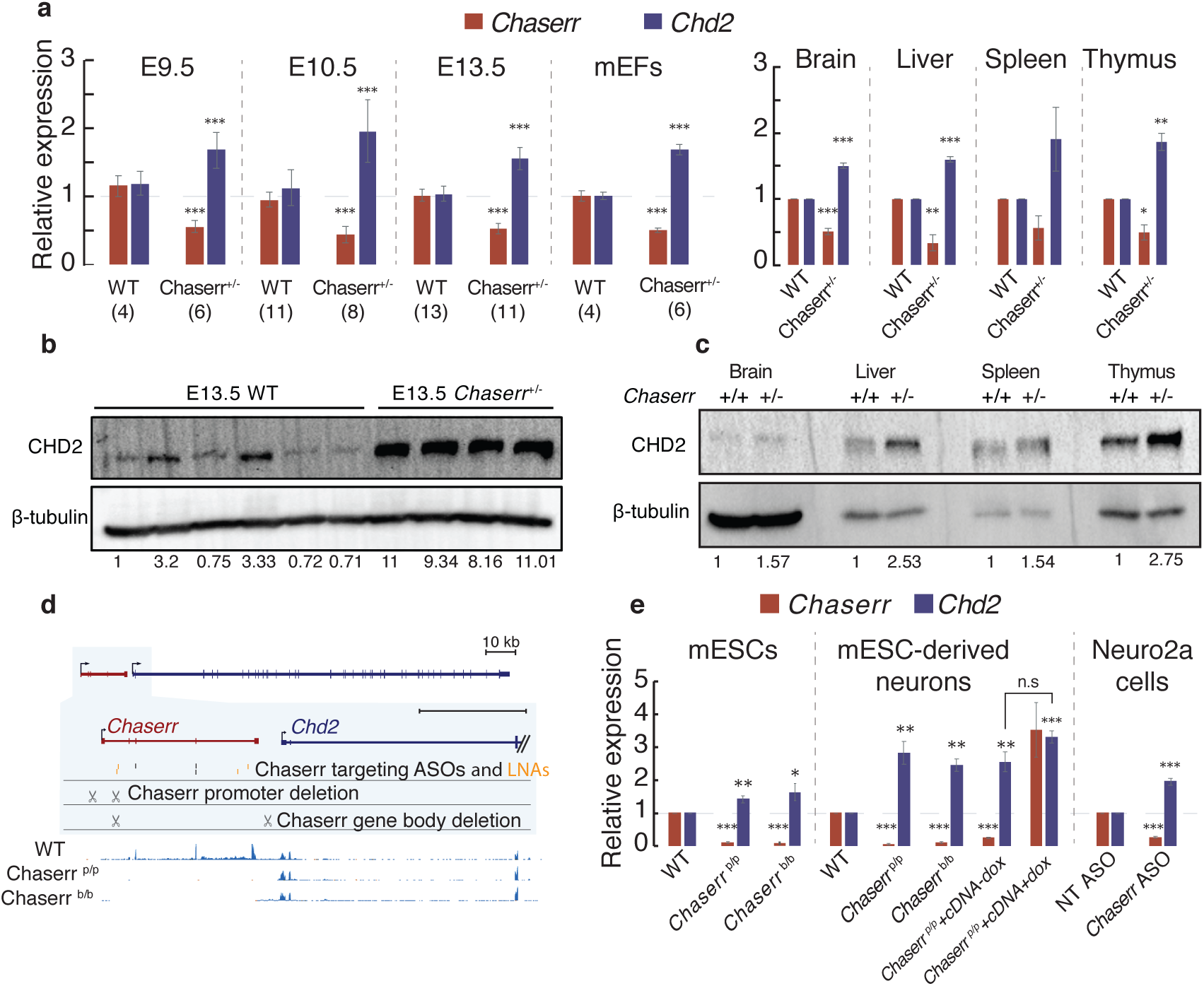
*Chaserr* represses *Chd2* expression. **a,** qPCR of the indicated genes in whole embryos from the indicated developmental stage, mEFs, or adult tissues with the indicated background. n for each group of embryos or mEFs is indicated in parentheses. Normalized to *Actb*. n=3 for the tissues. **b**, Western blot for the indicated protein in individual E13.5 whole embryos from indicated background. **c,** as in b for the adult tissues. **d,** Top: Scheme of regions targeted by gRNAs used to generate *Chaserr*^p/p^, *Chaserr*^b/b^ mESCs, or the antisense reagents; Bottom: RNA-seq read coverage in mESCs from the indicated background. **e,** qRT-PCR of the indicated genes in the indicated backgrounds or treatments. Normalized to *Actb*. n≥3. Error bars show S.E.M. *P<0.05 **P<0.01 and ***P<0.001 (two-sided t-test).

As *Chaserr*^-/-^ mice could not be generated, in order to study regulation of Chd2 in *Chaserr*^−/−^ cells and efficiently compare different *Chaserr* perturbations we used CRISPR/Cas9 in mouse embryonic stem cell (mESCs) to engineer clones with either homozygous loss of *Chaserr* promoter and first exon (*Chaserr*^p/p^), or deletion of the rest of the *Chaserr* gene body from the first intron to just downstream of *Chaserr* polyadenylation site (*Chaserr*^b/b^) (**Fig. 2d**). Importantly, the *Chaserr*^b/b^ line has an intact *Chaserr* promoter, and thus helps address the possibility of a competition between the *Chaserr* and *Chd2* promoters. *Chaserr* expression was abolished in both mutant lines, as evident in RNA-seq data (**Fig. 2d**), leading to significant *Chd2* mRNA upregulation (**Fig. 2e**). This up-regulation is consistent with a previous study that examined *Chaserr* as part of a panel of lncRNAs (*Chaserr* was referred to as ‘linc2025’ in that study) and also observed an increase in *Chd2* upon deletion of *Chaserr* promoter^27^. *Chd2* was also significantly upregulated in neurons derived from *Chaserr*^p/p^ and *Chaserr*^b/b^ mESCs (**Fig. 2e**). Similarly, targeting of *Chaserr* in Neuro2a cells using antisense oligonucleotides (ASOs) or LNA Gapmers reduced *Chaserr* expression by ~75%, and led to a significant increase in *Chd2* mRNA and protein levels (**Fig. 2e** and **Extended Data Fig. 2a-b**). We note that other perturbation methods, like insertion of polyA sites or ribozymes^28^ are not effective for perturbing *Chaserr* (^27^ and data not shown). We conclude that an intact *Chaserr* represses *Chd2* expression in all the systems we studied.

In order to test whether the mature *Chaserr* RNA is sufficient for repression of *Chd2*, we infected the *Chaserr*^p/p^ mESCs with a lentivirus carrying a doxycycline (dox)-inducible *Chaserr* cDNA. *Chaserr* was overexpressed relative to the WT levels by ~3.5-fold following Dox addition, but *Chd2* mRNA expression was not affected compared to no Dox control (**Fig. 2e**). We conclude that either the RNA product of *Chaserr*, or transcription from its endogenous locus (which might be affected by the ASO/Gapmer cleavage of the nascent transcript), are required for repression of *Chd2* mRNA.

To evaluate if the broader genomic context is important for *Chaserr* function, we reconstituted the *Chaserr* locus on a plasmid, which contains a fragment of the mouse genome from the *Chaserr* promoter to the start codon of *Chd2*, excluding *Chaserr* introns, cloned immediately upstream of a firefly luciferase coding sequence (**Extended Data Fig. 2c**). Deletion of *Chaserr* promoter in this system led to a ~50% increase in luciferase expression (**Extended Data Fig. 2c**), consistent with an increase we observed in the endogenous setting. This suggests that the information required for at least part of *Chaserr* activity is encoded in the tested locus.

We next examined whether the early lethality of mice with *Chaserr* loss-of-function is mediated by increased *Chd2* levels. We used CRISPR/Cas9 with a single gRNA targeted to the 3^rd^ exon of *Chd2* (**Fig. 1d** and **Extended Data Fig. 2d**) to generate *Chd2*^m/m^ mice that had a two bp deletion in the coding sequence of *Chd2*. This mutation had a mild effect on Chd2 mRNA expression, but dramatically reduced CHD2 protein levels, as expected from a frameshift in the main coding frame, with the residual protein possibly emanating from an alternative start codon (**Fig. 3a**, note that the monoclonal antibody we used recognizes a peptide in the C-terminal part of CHD2). *Chd2*^m/m^ mice were born at expected Mendelian ratios (**Extended Data Fig. 2e**) and had no gross developmental phenotypes, similarly to the *Chd2*^tm1b(EUCOMM)Hmgu^ mice generated by insertion of a LacZ cassette into the 2^nd^ intron and deletion of 3^rd^ exon of *Chd2^29^*. This enabled us to breed *Chaserr*^+/–^ mice with *Chd2*^m/m^ mice (**Fig. 3b**). We intercrossed the resulting *Chaserr*^+/–^ *Chd2* ^m/+^ offspring, and out of 44 pups born from such crosses, only 14 (~32%) were *Chaserr^+/–^* (**Extended Data Fig. 2f**), suggesting that one hypomorphic allele of *Chd2* is not sufficient for compensating for loss of one *Chaserr* allele. Indeed, CHD2 protein levels were substantially higher in *Chaserr*^+/–^ *Chd2*^m/+^ mice when compared to their *Chaserr*^+/+^ *Chd2*^m/+^ littermates (**Fig. 3c**), potentially explaining the neonatal lethality and growth retardation (**Extended Data Fig. 2g**).

**Figure 3.**
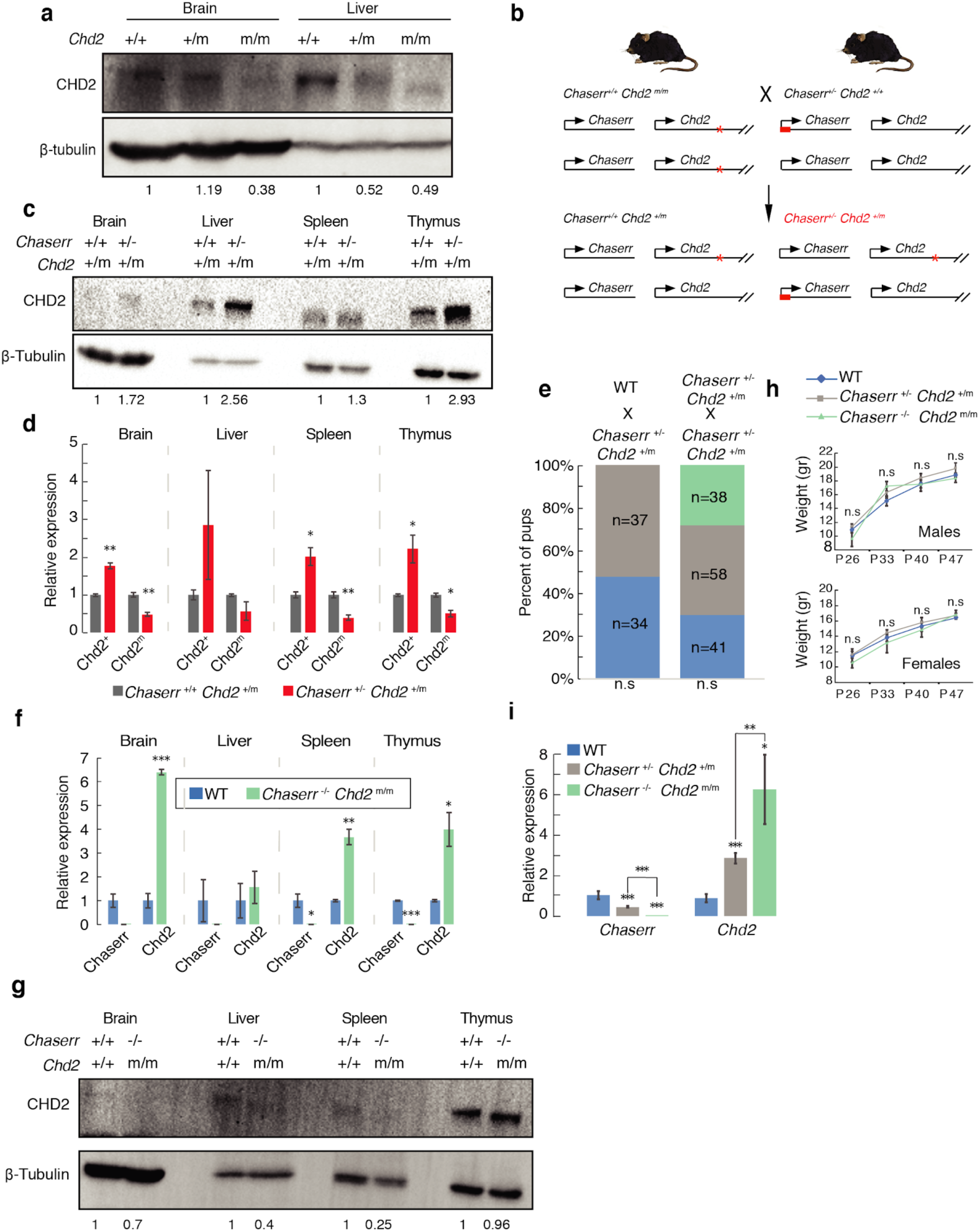
*Chaserr* regulates *Chd2* expression in *cis*. **a,** CHD2 western blot in the indicated tissues from mice with the indicated genotypes. **b,** Scheme of the cross between *Chaserr*^+/–^ and *Chd2*^m/m^ mice. **c,** Western blot in the indicated tissues from mice with the indicated genotypes. **d,** *Chd2* allele-specific expression in the indicated backgrounds in four different mouse tissues. Normalized to *Actb*. n=3 for the indicated tissues. **e,** Neonate survival rates for the *Chaserr*^+/–^ *Chd2*^m/+^ intercross. **f,** qRT-PCR in the indicated adult tissues and backgrounds. Normalized to *Actb*. n=3 for the indicated tissues **g,** CHD2 western blot from the indicated tissues and backgrounds. **h,** Pup weights for the *Chaserr*^+/–^ *Chd2*^m/+^ intercross. n=5–19 pups per genotype per time point. **i,** qRT-PCR for mEFs with the indicated background. Normalized to *Actb*. WT n=5, *Chaserr*^+/-^ *Chd2*^+/m^ n=20, *Chaserr*^+/-^ *Chd2*^+/m^ n=5. Error bars show S.E.M. *P<0.05 **P<0.01 and ***P<0.001 (two-sided t-test).

The *Chaserr*^+/–^ *Chd2*^m/+^ offspring that survived allowed us to test with allele-specific qRT-PCR whether *Chaserr* affected *Chd2* expression from the *cis* allele *in vivo* (**Extended Data Fig. 2h**). Only *Chd2* from the *Chaserr*^−^ allele was up-regulated in *Chaserr*^+/−^ *Chd2*^m/+^ mice when compared to *Chaserr*^+/+^ *Chd2*^m/+^ littermates (**Fig. 3d**), suggesting that *Chaserr* represses *Chd2* through *cis*-acting regulation. Interestingly, the *Chd2* from the *Chaserr*^+^ allele was repressed compared to WT levels, hinting at a possible feedback regulation resulting from excess CHD2 expression (see Discussion).

In order to directly test whether the severe *Chaserr* loss-of-function phenotype is mediated by CHD2 over-expression, we used CRISPR/Cas9 to delete the *Chaserr* promoter and first exon in a *Chd2*^m/m^ mouse (**Extended Data Fig. 3a**), and thus generated a model in which loss of *Chaserr* increases the expression of a hypomorphic allele of *Chd2*. Out of 137 pups born from intercrosses of *Chaserr*^+/–^ *Chd2*^+/m^ mice, 38 were *Chaserr*^−/−^ *Chd2*^m/m^ (~27%), 58 *Chaserr*^+/–^ *Chd2*^+/m^(~42%), and 41 WT (~30%), which did not deviate significantly from normal Mendelian ratios (P =0.2; **Fig. 3e** and **Extended Data Fig. 3b,c**), despite up to 6-fold up-regulation of *Chd2* mRNA in different tissues of *Chaserr*^−/−^ *Chd2*^m/m^ mice (**Fig. 3f**). Despite the mRNA up-regulation, CHD2 mutant protein levels in tissues from *Chaserr*^−/−^ *Chd2*^m/m^ mice were not higher than the WT protein levels in WT mice (**Fig. 3g**). *Chaserr*^−/−^ *Chd2*^m/m^ and *Chaserr*^+/–^ *Chd2*^+/m^ mice showed no significant differences in weight (**Fig. 3h** and **Extended Data Fig. 3d**), and no obvious phenotypes, in stark contrast to the common and pleiotropic phenotypes observed in *Chaserr*^+/–^ neonates on *Chd2*^+/+^ background. A hypomorphic allele of *Chd2* can thus rescue the lethality caused by loss of *Chaserr*, when the two occur on the same allele.

To further characterize the changes in levels of *Chd2* mRNA in the double mutants, we isolated mEFs from E13.5 embryos from *Chaserr*^+/+^, *Chaserr*^+/–^, and *Chaserr*^−/−^ genotypes on *Chd2*^m/m^ background. qRT-PCR analysis showed that *Chaserr* expression is completely abolished in *Chaserr*^−/−^ *Chd2*^m/m^ mice, and that *Chd2* mRNA was significantly overexpressed in a manner that was inversely dependent on *Chaserr* dosage (**Fig. 3i**).

Having established that the severe phenotype resulting from *Chaserr* loss is mediated by CHD2, we were next interested to understand the consequences on gene expression of *Chaserr* loss and CHD2 up-regulation. We isolated *Chaserr*^−/−^ mEFs, which showed a >2-fold increase in Chd2 mRNA and protein levels (**Fig. 4a**), and used RNA-seq to characterize gene expression in *Chaserr*^+/+^ and *Chaserr*^+/–^ E9.5 and E13.5 embryos, adult brains, as well as *Chaserr*^+/+^, *Chaserr*^+/–^, *Chaserr*^−/−^, and *Chaserr*^−/−^ *Chd2*^m/m^ mEFs (**Fig. 4b**). As expected from the difference in the severity of the phenotype, we observed limited differential expression in *Chaserr*^+/–^ embryos or mEFs compared to WT, whereas substantial numbers of differentially expressed genes were found in *Chaserr*^−/−^ mEFs. The 1,493 significantly down-regulated genes (down by at least 25%, adjusted P<0.05) were enriched for GO categories such as nervous and connective tissue development and transcriptional regulation, and were very significantly enriched for genes whose loss-of-function in mice is associated with phenotypes such as “decreased length of long bones”, “decreased body weight”, “respiratory distress”, and “premature death” (**Supplementary Table 1**), closely related to the phenotypes observed in our *Chaserr* loss-of-function model. Specifically, some of the most down-regulated genes, such as *Ctsk*, *Cpt1c*, and *Dapk3*, which were also validated by qRT-PCR (**Fig. 4c**), are known to be required for proper embryonic development^30^.

**Figure 4.**
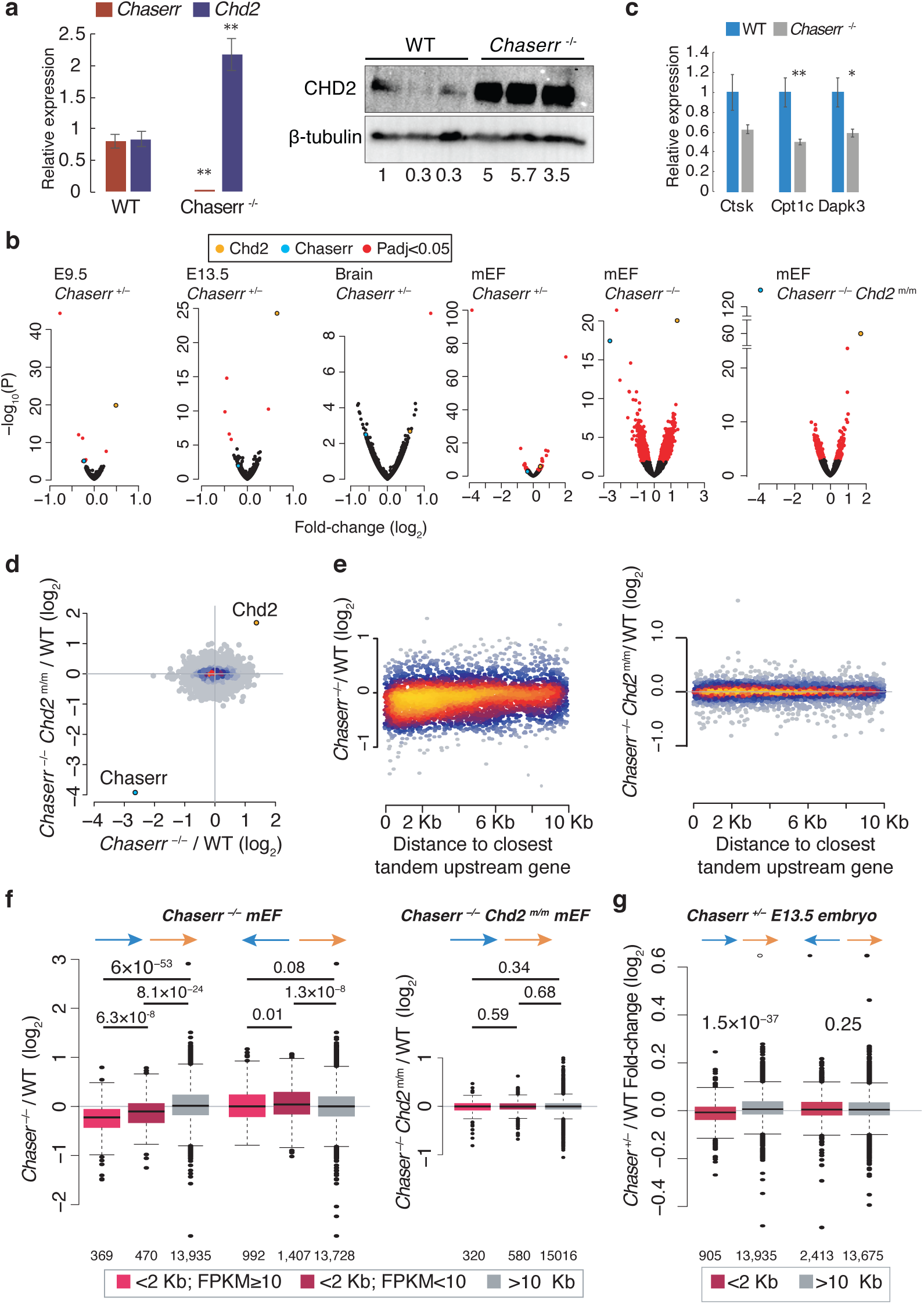
Transcriptional dysregulation following loss of *Chaserr*. **a,** qRT-PCR (left, WT n=4, *Chaserr*^−/−^ n=6) and Western blot (right) of *Chd2* expression in mEFs from the indicated background. n=3. qRT-PCR normalized to *Actb*. Western blot shows mEFs from three independent embryos for each background. **b,** Volcano plots for the indicated differential expression comparison using DESeq2 ^53^. **c,** qRT-PCR of the indicated genes in WT and *Chaserr*^−/−^ mEFs. Normalized to *Actb*. WT n=4, *Chaserr*^−/−^ n=6. **d,** Changes in gene expression in *Chaserr*^−/−^ and *Chaserr*^−/−^ *Chd2*^m/m^ MEFs. **e,** Change in expression in *Chaserr*^−/−^ or *Chaserr*^−/−^ *Chd2*^m/m^ vs. WT mEFs and distance to the 3’ end of the closest upstream gene transcribed from the same strand, for genes for which this distance is <10 Kb. **f,** Changes in gene expression in *Chaserr*^−/−^ or *Chaserr*^−/−^ *Chd2*^m/m^ vs. WT mEFs for genes with the indicated distances from the closest upstream gene. Genes with distance <2 Kb were further subdivided based on the expression levels of the upstream gene. **g,** As in e, for the E13.5 whole embryo RNA-seq. The number of genes in each group is indicated below the distance threshold. Error bars show S.E.M. *P<0.05, **P<0.01 (two-sided t-test).

In contrast, the 616 significantly up-regulated genes (up by at least 25%, adjusted P<0.05) were enriched for various RNA-related GO categories (**Supplementary Table 1**). Notably, changes in E13.5 embryo were significantly anti-correlated with those in E13.5 medial ganglionic eminence (MGE) of *Chd2*^+/–^ mice^22^ (Spearman R=–0.27, P<10^−100^). Milder anti-correlation was observed between dysregulation in the *Chaserr*^+/–^ adult brain and the *Chd2*^+/–^ adult hippocampus^22^ (R= –0.11, *P*= 8.8×10^−47^). Hundreds of genes were dysregulated in the *Chaserr*^−/−^ *Chd2*^m/m^ mEFs compared to WT mEFs from the same pregnancies, and these changes are likely due to *Chd2* loss-of-function, as there was no correlation between these changes and the changes in *Chaserr*^−/−^ mEFs (Spearman R=−0.003 P=0.68, **Fig. 4d**). These observations further support the notion that the downstream effects of loss of *Chaserr* are driven by up-regulation of *Chd2*.

Inspection of the loci of some of the most affected genes, such as *Ctsk* and *Dapk3* (**Fig. 4c**), led us to suspect that promoters of down-regulated genes might be preferentially found in close proximity to transcription termination sites (TTSs) of genes transcribed on the same strand (and thus potentially susceptible to transcriptional interference), in a similar organization to the *Chaserr*-*Chd2* locus. Indeed, we found a mild yet highly significant correlation between changes in gene expression in *Chaserr*^−/−^ mEFs and the distance to the TTS of the closest tandem upstream gene (Spearman R=0.2 P=2×10^−151^), with the effect observed mostly when the distance was shorter than ~6 Kb (**Fig. 4e**). In contrast, no such effect was observed in *Chaserr*^−/−^ *Chd2*^m/m^ mEFs (R=0.0055 P=0.45, **Fig. 4e**). Further, down-regulation was stronger when the expression of those upstream neighboring genes was higher (Spearman R=−0.17 P=3.4×10^−7^ between the change in expression of the downstream gene and the absolute expression of the upstream gene for intergenic distances <2Kb). Genes with a close and abundant upstream neighbor were significantly repressed in *Chaserr*^−/−^ mEFs, and in stark contrast, much smaller differences, and in opposite direction, were observed for neighboring genes transcribed on opposite strands, when considering distances between 5’ ends (**Fig. 4f**). Notably, in most cases of repression of close tandem downstream gene, both the upstream and the downstream gene were protein-coding (190 of the 194 pairs with intergenic distance <2 Kb and a reduction of at least 25% in the expression of the downstream gene in *Chaserr*^−/−^ mEFs). Similar trends were also found in data from E13.5 embryos, with much smaller effect sizes (**Fig. 4g**).

To further characterize the regulatory dysregulation, we performed ATAC-seq^31^ using WT and *Chaserr*^−/−^ mEFs. Consistently with the increase in *Chd2* production, we observed a mild increase of ~7% in accessibility of *Chd2* promoter in *Chaserr*^−/−^ mEFs compared to WT. In order to analyze the changes in accessibility throughout the genome following *Chaserr* depletion, we merged accessibility peaks from all samples, assigned them to the nearest gene, and classified the peaks as “Core promoter” (<300 nt from TSS), “Extended promoter” (>300nt and <2 kb from TSS), “Gene body”, or “Intergenic”, based on RefSeq annotations. Comparing WT and *Chaserr*^−/−^ mEFs, we found 84 and 348 regions with reduced or increased accessibility, respectively (t-test P<0.05), with changes observed in gene bodies and in promoters (**Fig. 5a**). Promoter peaks whose accessibility was significantly decreased in *Chaserr*^−/−^ mEFs were associated with concordant changes in gene expression (**Fig. 5b**). Strikingly, these peaks were predominantly found at promoters of genes separated by short intergenic regions from TTSs of other genes with the same relative orientation (**Fig. 5c**, median distance of 2.6 Kb compared to 43.9 Kb for unaffected promoters). All nine genes with significantly reduced core promoter accessibility, a tandem upstream neighbor within <5 Kb, and robust expression in the RNA-seq data were repressed by at least 25% (all with adjusted P<0.05, **Fig. 5d,f**). Notably, in these cases, the upstream gene was expressed at dramatically higher levels than the affected downstream gene (15-fold on average, **Fig. 5e**), and the expression of the upstream genes was not substantially affected by loss of *Chaserr* (**Fig. 5f**).

**Figure 5.**
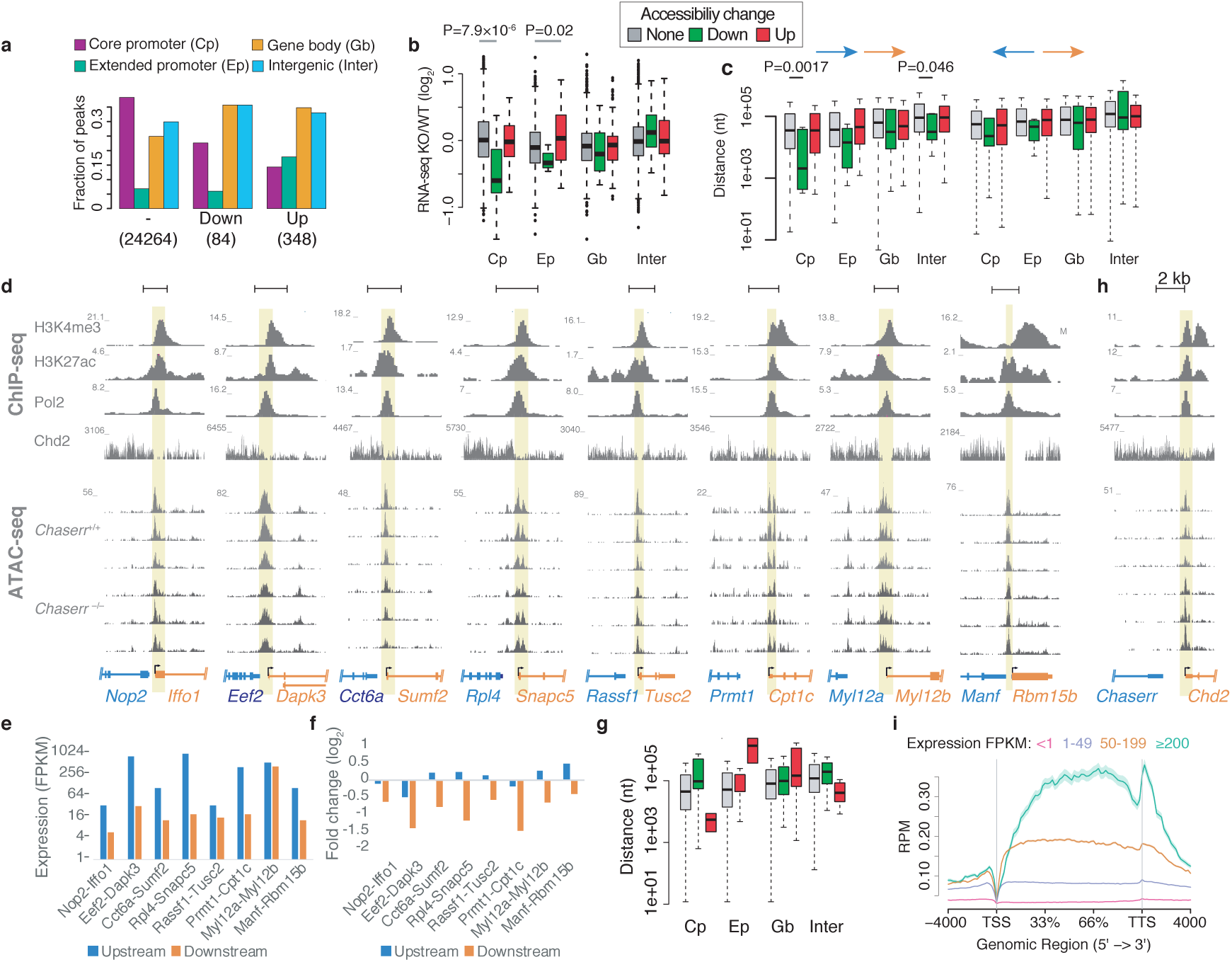
Changes in chromatin accessibility following loss of *Chaserr*. **a,** Distribution of ATAC-seq peaks that had reduced (‘Down’) or increased (‘Up’) accessibility in *Chaserr*^−/−^ mEFs among the indicated four groups. n for each group indicated in parentheses. **b,** Changes in gene expression of genes assigned to peaks in the indicated group (from a). Group names are as in legend of a. P values computed using two sided Wilcoxon rank-sum test. **c,** Distance from the closest gene on the same (left) or opposite (right) strand for the genes in the indicated group (from a). **d,** Loci of eight genes with reduced accessibility of peaks in their core promoters in the *Chaserr*^−/−^ background. Top: ChIP-seq coverage in mEFs from the ENCODE project. Middle: CHD2 MNase-ChIP-seq read coverage in mESCs^13^. Bottom - read coverage in ATAC-seq data from mEFs isolated from three embryos from each background. **e,** RNA-seq expression levels for the indicated pairs of genes in WT cells. **f,** Differential expression in *Chaserr*^−/−^ mEFs compared to WT mEFs for the indicated gene pairs. **g,** Same as c left, but using data from *Chaserr*^−/−^ *Chd2*^m/m^ mEFs, compared to WT mEFs. **h,** As in d, for the *Chaserr*/*Chd2* locus. **i,** Metagene of CHD2 occupancy^13^ on genes in mESCs, divided into four groups based on their expression FPKM.

To test if these effects stem from CHD2 over-expression, we used the same methodology to compare *Chaserr*^−/−^ *Chd2*^m/m^ mEFs with WT mEFs matched from the same pregnancies, and observed fewer changes in accessibility (67 and 26 peaks with decreased and increased accessibility, respectively, based on the same criteria as above), and no preferential association of decreased promoter accessibility with distance to tandem upstream genes (**Fig. 5g**), suggesting that the transcriptional interference in *Chaserr*^−/−^ cells is driven by excess CHD2 levels.

The presence of a TTS closely upstream of the promoters most affected by loss of *Chaserr* (**Fig. 5d**) resembles the organization of the *Chaserr* and *Chd2* locus (**Fig. 5h**). We observed substantial occupancy of CHD2 protein (using tagged CHD2 MNase-ChIP-seq data from mESCs^13^) in the short intergenic regions between the upstream TTS and the promoters of the repressed genes, as between *Chaserr* TTS and *Chd2* (**Fig. 5d,h**), suggesting that the effect seen on the accessibility of these promoters and gene expression might be a direct result of increased CHD2 activity in the intergenic regions. Concordantly with these individual cases, a metagene analysis of tagged CHD2 occupancy across the gene body showed that CHD2 preferentially associates with ~2Kb regions immediately downstream of TTSs (**Fig. 5i**). CHD2 thus appears to function in regions downstream of TTSs in WT cells, and the CHD2 hyperactivity caused by loss of *Chaserr* leads to repression of promoters adjacent to those regions.

## Discussion

Decoding lncRNA functions is a formidable challenge, in particular because of the substantial heterogeneity in their biology and modes of action. A prominent group of vertebrate lncRNAs, including some of the most conserved ones, are found in regions flanking genes involved in transcription, including numerous chromatin modifiers. We hypothesized that such co-localization may imply a connection between the biology of the chromatin modifier and the mode of action of the lncRNA, potentially through a feedback loop. By focusing on one such pair that is particularly highly conserved in evolution, we uncovered a lncRNA-mediated circuit that regulates the expression of *Chd2*.

Negative autoregulatory feedback loops are prevalent in RNA biology, and help tune levels of factors involved in mRNA splicing, polyadenylation, editing, and modifications, and in processing of small RNAs^32–38^. There are also examples of such feedback loops regulating transcription-related genes^39^, but to the best of our knowledge, there are no known autoregulatory loops involving chromatin modifiers in mammals. We propose a model in which the *Chaserr*-*Chd2* tandem organization forms a negative feedback loop that tunes CHD2 levels. The genomic occupancy of CHD2 in regions immediately downstream of TTSs (**Fig. 5i**), as well as the enrichment of CHD2-mediated accessibility changes in gene bodies (**Fig. 5a**), suggest that CHD2 acts in TTS-proximal regions. Accordingly, we show that under conditions of excess levels of CHD2 protein, this TTS-proximal occupancy is associated with transcriptional interference on downstream neighbors of highly-expressed genes, especially when the intergenic regions between them are particularly short (**Fig. 4e-g**, **Fig. 5c-g**). Due to the short intergenic region between *Chaserr* and *Chd2*, we suggest that such transcriptional interference serves as the basis of a negative feedback loop that leads to repression of *Chd2* production when CHD2 protein levels are high, thus maintaining a tight control on its expression. In cells with *Chaserr* loss-of-function, this loop is compromised, and CHD2 is upregulated. Supporting this model, in *Chaserr*^+/–^ *Chd2*^m/+^ tissues, increase in the CHD2 protein leads to repression of transcription of the *Chd2*^m^ allele, which is found next to an intact *Chaserr* (**Fig. 3d**).

The exact mechanism by which *Chaserr* represses *Chd2* expression remains unknown. The simplest feedback model suggests that *Chaserr* transcription, or perhaps its termination and cleavage and polyadenylation, are important. Such a mechanism would resemble the activity of the SRG1 noncoding RNA in the yeast *S. cerevisiae*, that represses the transcription of SER3, which is found immediately downstream of SRG1 and on the same strand, by deposition of nucleosomes at the SER3 promoter that prevent binding of transcription activators^40–44^. The genomic arrangement of SRG1 and SER3 resembles that of *Chaserr* and *Chd2*, though the 3’ end of SRG1 overlaps SER3 promoter^41^, and the distance between their promoters is much shorter (~500 bp between SRG1 TSS and SER3 TSS vs. ~17 kb between the TSS of *Chaserr* and TSS of *Chd2*). However, there is evidence that the RNA product of *Chaserr* is important for its function. The exon-intron architecture and sequence of *Chaserr* are highly conserved, suggesting that production of a particular RNA species, or splicing at particular locations, is important either because of the RNA product itself, or because they modulate the amount of transcription or Pol2 velocity upon encounter with the ~2 Kb intergenic region between *Chaserr* and *Chd2*. The similar effects we observed when deleting *Chaserr* promoter or gene body, or when targeting the RNA with ASOs or Gapmers, also point to the importance of the RNA product, though we cannot rule out that ASOs and Gapmers may affect transcription by cleaving the nascent transcript.

Both *Chaserr* and *Chd2* appear to be under the shared control of several enhancer regions found in the ~200 Kb gene desert upstream of *Chaserr* (**Extended Data Fig. 4**), and those enhancers potentially regulate both genes together, which can explain the tight co-expression between *Chaserr* and *Chd2* mRNA (**Fig. 1c**). We propose that shared regulation of both genes is potentially important for maintaining the integrity of the feedback mechanism, as we found that expression levels of the upstream gene appear to be consequential for transcriptional interference resulting from CHD2 hyperactivity (**Fig. 4f**).

Interestingly, in contrast to previous reports based on a gene trap inserted in intron 27 of *Chd2^17,45,46^*, nearly complete loss of CHD2 protein appears to be largely compatible with normal viability in mice in our hands, and in the similar mouse model generated by the EUCOMM consortium^18^. The previous observations of embryonic lethality of CHD2 loss-of-function can potentially be attributed to a dominant negative effect of the truncated protein product created by the gene trap. In contrast to the CHD2 loss models, we report here that increase in CHD2 levels brought about by loss of even a single copy of *Chaserr* is toxic and leads to perinatal lethality. We note that this is one of the most severe phenotypes reported so far for loss of a lncRNA^6,47^. The severe, CHD2-mediated phenotype is consistent with the high conservation of *Chaserr*-*Chd2* genomic organization and sequence during 500 million years of vertebrate evolution. Of the lncRNAs annotated in human and mouse, only ~5% (~100 genes) have evidence of conservation in fish^26,48^, and we were able to identify homologs of *Chaserr* in every vertebrate species we examined (**Fig. 1a**).

Our results have important implications from the therapeutic perspective. Individuals that bear mutations in the *CHD2* gene exhibit epilepsy and neurodevelopmental disorders^19^. In all described cases, these individuals are haploinsufficient for *CHD2*, and so bear an intact WT copy of CHD2. Therefore, increase of CHD2 expression through perturbation of *Chaserr*, e.g., using antisense oligonucleotides, might have a therapeutic benefit. Importantly, *Chaserr* is highly conserved between human and mouse (**Fig. 1a**), and targeting of the human *Chaserr* using Gapmers in human MCF7 and SH-SY5Y cells leads to an increase in *CHD2* mRNA and protein levels (**Extended Data Fig. 5**), suggesting that the results we observe in the mouse model are of direct relevance to human *Chaserr*.

Similarly to many other chromatin remodelling factors, that are increasingly implicated in human disease, the precise molecular function of *Chd2* and the direct consequences of its dysregulation are poorly understood. Here, we could leverage the similarity between the genomic arrangements of *Chaserr* and *Chd2* and those of the genes most affected by *Chaserr* perturbation to highlight the potential role of chromatin remodelling that CHD2 plays downstream of TTSs. Decoding lncRNA functions can therefore provide important insight into other layers of gene regulation. As chromatin modifiers are often flanked by lncRNAs, some as highly conserved as *Chaserr*, we expect that further research into the functions of these lncRNAs may uncover new paradigms in chromatin biology.

## Materials and Methods

### Animals

The study was conducted in accordance with the guidelines of the Weizmann Institutional Animal Care and Use Committee (IACUC). C57black6 Ola HSD mice were purchased from Harlan Laboratories (Rehovot, Israel). All other mouse strains were bred and maintained at the Veterinary Resources Department of the Weizmann Institute.

### Generation of KO mice

The CRISPR KO mice were generated as in (Wang et al., 2013). All KO mice were generated by standard procedures at the Weizmann transgenic core facility. To generate *Chaserr* KO mice we used four single guide RNAs (sgRNAs 1-4, see **Supplementary Table 2**), two targeting regions before the *Chaserr* transcription start site and two targeting regions in the first intron, which resulted in two founder lines. *Chaserr* mutant mice were genotyped using primers flanking the targeted region followed by Sanger sequencing. To generate *Chd2* mutant mice we used one sgRNA targeted to the third exon of Chd2 mRNA. Chd2 mutant mice were genotyped by amplicon sequencing which identified a founder with an ORF shift. Amplicon library preparation was as follows: Chd2 mutant locus was amplified using Chd2 genotype nesting primers (**Supplementary Table 2**). Thereafter 1µl PCR reaction was used as template for the addition of R1 and R2 Illumina adaptors (R1/R2_Exon3_Chd2, **Supplementary Table 2**), followed by fragment AMPure (Beckman Coulter, A63881) size selection and cleanup. A second PCR reaction was used in order to add sequencing barcodes to the amplicons, followed by AMPure size selection. Indexed amplicons were pooled and sequenced on NextSeq 500. To genotype *Chd2^m^* lines routinely, we used PCR with primers flanking the mutated region and digested the PCR product with the DdeI restriction enzyme whose recognition sequence is compromised in the mutated DNA. The fragment sizes were then analyzed on a 2% agarose gel. To generate *Chaserr^−^*^/–^ *Chd2*^m/m^ mice we used *Chd2*^m/m^ background and continued as described above for generation of the *Chaser^−^* allele, using only two sgRNAs (sgRNA2+3, **Supplementary Table 2**). sgRNA injection were done on CB6F1 Ola HSD mice which were later backcrossed with C57BL/6 Ola HSD. All the experiments were done on 4 to 15-week-old mice from F2–F5 generations and E9.5–E18.5 developmental time points.

### Tissue culture

mESCs were routinely cultured in mouse ES medium (mESM) consisting of: 500ml DMEM (Gibco, 11965-092), 15% ES-grade Fetal Calf Serum (Biological Industries), sodium pyruvate 1mM (Gibco, 11360-039), nonessential amino acids 1x (Gibco, 11140-035), 0.1mM b-mercaptoethanol (Sigma, M6250-250ML), penicillin-streptomycin (Biological Industries), and 1000U/ml LIF (Weizmann Proteomics Unit).

All other cell lines were routinely cultured in DMEM containing 10% fetal bovine serum and 100 U penicillin/0.1 mg ml^−1^ streptomycin at 37° C in a humidified incubator with 5% CO2. Cell lines were routinely tested for mycoplasma contamination.

Primary mEFs were isolated as previously described^49^.

### Neuronal differentiation

Neuronal differentiation was performed as previously described (Ying et al., 2003). mESCs were first grown in the absence of mEFs for two passages and then seeded on gelatin-coated plates at a density of 1.5×10^5^ in N2B27 medium: 1:1 mixture of DMEM/F12 (Sigma) supplemented with N2 (Gibco), and Neurobasal medium (Gibco) supplemented with B27 (Gibco), 1X Glutamax (Gibco), 0.1 mM β-mercaptoethanol (Sigma), 100 U/ml penicillin, and 0.1 mg/ml streptomycin (Biological industries). After four days under these conditions, 3×10^5^ cells were re-plated on Poly-D-Lysine (Sigma, P6407) and Laminin (Life, 23017-015) coated plates, in N2B27 medium supplemented with 20ng/ml FGF2 (Peprotech, 100-18B-50/100-18B-100). After 24 hr, FGF2 was removed and cells were cultured for three additional days.

### Transfections

mESCs were transfected with electroporation the Lonza protocol (http://bio.lonza.com/fileadmin/groups/marketing/Downloads/Protocols/Generated/Optimized_Protocol_309.pdf). HEK293T cells were transfected with were performed using PolyEthylene Imine (PEI)^50^ (PEI linear, M*r* 25,000, Polyscience). Neuro2A transfection: 2×10^5^ were seeded in 6 well-plate and transfected using Lipofectamine 3000 (Life, L3000-008) with LNA1, LNA2 or mix of LNA1-4 (LNA KD) or with ASO1, ASO2, ASO3 or mix of ASO1-3 (ASO KD) to final concentration of 50 nM. For transfection of MCF7 cells, 2×10^5^ were seeded in a 6-well plate and transfected using PEI (as previously described) with LNA h1 and/or LNA h2 to final concentration of 50 nM. For luciferase experiments, 20×10^3^ cells were seeded in a 24-well plate and co-transfected using PEI with pGL3-Chaserr-reporter vector (100 ng) and psiCHECK-1 (100 ng, Promega, C8011). Firefly and Renilla expression were measured using Dual-Luciferase Reporter Assay System (Promega, E1910) following manufacturer’s protocol. For transfection of SH-SY5Y cells, 2×10^5^ cells were seeded in a 6-well plate and transfected using DharmaFECT 4 Transfection Reagent (Dharmacon, T-2002-03) following manufacturer’s protocol with LNA h1 and/or LNA h2 to final concentration of 50 nM. End time point for all KD experiments and luciferase assay was at 48 hr post transfection.

### Genome editing in mESCs

To generate mESC *Chaserr*^p/p^ mESCs, 2×10^6^ mEF-depleted cells were transfected with sgRNAs 2+3 and pCas9_GFP (a gift from Kiran Musunuru, Addgene #44719). To generate *Chaserr*^b/b^ mESCs, 2×10^6^ cells were transfected with sgRNAs 2+5 and pCas9_GFP. The next day fresh mESC medium was supplemented with 1µg/ml puromycin (Invivogen, ant-pr-1) for 72 hr, while replacing medium each 24 hr. Next, 3–4×10^3^ cells were seeded at low density on 10 cm plate until single colonies formed, thereafter colonies were picked, expanded, genomic deletion was verified by PCR, sequencing, and expression was tested by RT-qPCR.

### RNA and RT-qPCR

Total RNA was extracted from different cell lines and mouse tissues, using TRIREAGENT (MRC) according to the manufacturer’s protocol. cDNA was synthesized using qScript Flex cDNA synthesis kit (95049, Quanta). Fast SYBR Green master mix (4385614) was used for qPCR.

### RNA-seq

Strand-specific mRNA-seq libraries were prepared from 500–4000 ng total RNA using the TruSeq Stranded mRNA (Illumina) or SENSE mRNA-Seq (Lexogen) library preparation kits, according to the manufacturer protocols. Libraries were sequenced on a NextSeq 500 to obtain 38–50 nt paired-end reads. Coverage tracks for the UCSC genome browser were prepared by aligning reads to the mm9 genome assembly using STAR^51^. Gene expression levels were quantified using RSEM^52^ and a RefSeq gene annotation database which was manually edited to correct the annotation of the last exon of *Chaserr*. Differential expression was computed using DESeq2 with default settings^53^. Genomic context was also analyzed using the RefSeq gene annotations. RNA-seq and ATAC-seq datasets are deposited in GEO database under the accession GSE124375 (reviewer token wtkleowgbhotjgn). RNA-seq data from previous studies were downloaded from the SRA database, and quantified using RSEM with the same annotation file.

### Western Blot

Total protein was extracted from tissues and cell lines by lysis with RIPA supplemented with protease inhibitors and DTT 1mM. Proteins were resolved on 8-10% SDS-PAGE gels and transferred to a Polyvinylidene difluoride (PVDF) membrane. After blocking with 5% non-fat milk in PBS with 0.1% Tween-20 (PBST), membranes were incubated with the primary antibody followed by the secondary antibody conjugated with horseradish peroxidase. Blots were quantified using Image Lab software. Primary antibodies as follows: anti-Chd2 (Millipore, MABE873), anti-β-tubulin (Sigma, T4026). Secondary antibodies as follows: anti-rat (AP136P), anti-mouse (115-035-003).

### ATAC-seq

ATAC-seq was performed as previously described (Buenrostro et al., 2013). 5×10^4^ mEFs derived from WT, *Chaserr^−^*^/–^, or *Chaserr*^−/−^ *Chd2*^m/m^ backgrounds. Libraries were sequenced with paired-end sequencing on Illumina NextSeq 500. Reads were aligned to the mm9 genome assembly using Bowtie2^54^ and peaks were called using MACS2^55^. Normalized read coverage files were computed by MACS2. Peaks form individual samples were merged using bedtools^56^ and read coverage in the peaks was computed using bigWigAverageOver bed UCSC utility^57^. Differential peaks were then called using two-sided t-test. Peaks were assigned to the closest RefSeq gene and annotated as “Core promoter” if they fell within 300 nt of a TSS; “Extended promoter if they fell within 2,000 nt of a TSS; “Gene body” is they overlapped a transcription unit; or “Intergenic” otherwise.

### Single-molecule FISH

Stellaris probe libraries targeting *Chaserr* introns or exons were designed using the Biosearch Technologies server (**Supplementary Table 3**) and ordered from Biosearch Technologies.

Duodenum sections were hybridized according to a previously published protocol ^58^ with the exception of using 25% formamide and 6µM sections. DAPI (Sigma-Aldrich, D9542) and a FITC-conjugated antibody against E-Cadherin (BD Biosciences, 612131) were used to identify the nucleus and cell-membrane, respectively. Imaging was performed on a Nikon-Ti-E inverted fluorescence microscope with a 100 × oil-immersion objective and a Photometrics Pixis 1024 CCD camera using MetaMorph software as previously described ^59^. The analysis was carried on ImageM, a custom Matlab program ^60^ to compute single-molecule mRNA concentrations in the nucleus or cytoplasm by segmenting each cell manually according to the cell borders and the nucleus.

### Cycloheximide treatment

mESCs and mEFs were treated with DMSO (vehicle) or CHX (Sigma #C7698) 100µg/mL and collected at the indicated time points for RNA/protein analysis.

### Extraction of cytoplasmic and nuclear RNA

Cells were washed twice with ice-cold PBS, then scraped with ice-cold buffer A (EGTA 15µM, EDTA 10µM, protease inhibitor cocktail (Sigma, P8340) and RNase inhibitor (ERX-E4210-01)) and centrifuged at 400g, 4°C for 5’ min. Supernatant was discarded, fresh buffer A was added and the pellet was mechanically pipetted with 21G followed by 27G syringe. Cells were then centrifuged at 2000g, 4°C for 5 min and the syringe step was repeated. The cells were then centrifuged at 2500g, 4°C for 5 min. The pellet was then kept as the nuclear fraction and the supernatant was centrifuged again at 6000g, 5 min followed by another supernatant collection (clean cytosolic fraction). Nuclear pellet was then washed three times with buffer A. RNA was extracted TRIREAGENT (MRC).

### Molecular cloning

Guide RNAs were designed using CHOPCHOP ^61^. Cloning of plasmids were done following Zhang Lab General Protocol (http://www.genome-engineering.org/crispr/wp-content/uploads/2014/05/CRISPR-Reagent-Description-Rev20140509.pdf) using phU6-gRNA^62^, (a gift from Charles Gersbach, Addgene plasmid # 53188) or pKLV-U6gRNA(BbsI)-PGKpuro2ABFP^63^ (a gift from Kosuke Yusa, Addgene plasmid # 50946). For luciferase experiments, a synthetic sequence containing *Chaserr* promoter, *Chaserr* cDNA, the intergenic region between *Chaserr* and *Chd2*, and *Chd2* promoter were synthesized by hylabs and cloned to into pGL3-basic backbone (Promega). To generate the plasmids with deleted fragments we used the following restriction enzymes followed by ligation: (i) *Chaserr* promoter deletion: XhoI+PvuII; (ii) *Chaserr* cDNA+intergenic deletion: PvuII+BglII; (iii) *Chaserr* cDNA+intergenic+Chd2 promoter: PvuII+HindIII; (iv) *Chaserr* promoter+*Chaserr* cDNA+intergenic: XhoI+BglII; (v) Chd2 promoter: BglII+HindIII.

### *Chaserr* cloning and lentiviral production

cDNA from Source BioSicence clone C130076G01 was amplified with PCR adding restriction sites for NheI at 5’ end and AgeI at 3’ end (**Supplementary Table 2**) and cloned into pLIX_402 vector (a gift from David Root, Addgene #41394) using restriction ligation. To produce viruses, HEK293T (2.5×10^6^, 10cm plate) cells were transfected using PEI with psPAX2 (3.5µg, a gift from Didier Trono, Addgene #12260), pMD2.G (1.5µg, a gift from Didier Trono, Addgene, #12259) and pLIX_402-Chaserr (5µg). Viruses were collected 48 hr post-transfection and filtered through 0.45µm sterile filters. Viruses were supplemented with polybrene (1:1000, Sigma, 107689-10G) upon cell infection.

### Micro-CT scanning and analysis

Prior to Micro-CT scanning the mice were anesthetized using IP injection of a mix of Xylazine (10mg/Kg) and Ketamine (100mg/Kg). Mice were scanned using a micro-CT device TomoScope ^®^ 30S Duo scanner equipped with two source-detector systems. The scanner uses two x-ray sources and a detector system that are mounted on a gantry that rotates around a bed holding the animal. The operation voltages of both tubes were 40kV. The integration time of protocols was 90ms (360 rotations) for 3cm length and axial images were obtained at an isotropic resolution of 80µm. Due to the maximum length limit, to cover the whole mouse body, imaging was performed in two-three parts with overlapping area and then all slices merged to one dataset representing the entire ROI. The radiation dose range was 0.9Gy. All micro-CT scans were reconstructed using a filtered back-projection algorithm using scanner software. Then the reconstructed data sets for each mice were merged to one data set using ImageJ software. 3D volume rendering images were produced using Amira Software.

